# Multitask learning for Transformers with application to large-scale single-cell transcriptomes

**DOI:** 10.1101/2020.02.05.935239

**Authors:** Minxing Pang, Jesper Tegnér

**Affiliations:** Nankai University; King Abdullah University of Science and Technology Karolinska Institutet

## Abstract

Recent progress in machine learning provides competitive methods for bioinformatics in many traditional topics, such as transcriptomes sequence and single-cell analysis. However, discovering biomedical correlation of cells that are present across large-scale data sets remains challenging. Our attention-based neural network module with 300 million parameters is able to capture biological knowledge in a data-driven way. The module contains high-quality embedding, taxonomy analysis and similarity measurement. We tested the model on Mouse Brain Atlas, which consists of 160,000 cells and 25,000 genes. Our module obtained some interesting findings that have been verified by biologists and got better performance when benchmarked against autoencoder and principal components analysis.

## 1 Introduction

Rapid progress in the development of single-cell RNA sequencing (scRNA-seq) technologies in recent years has provided many valuable insights into complex biological systems ([1],[2]). For example, scRNA-seq profiling of the adult mouse nervous system uncovers new cell classes and types across regions, providing a clearer picture of cell diversity by region and a reference atlas for studying the mammalian nervous system [3]. There are several machine learning algorithms developed to analyze the single-cell gene expression matrix [4], including visualization [5], imputation [6], data integration [7] and prediction of perturbation responses [8].

A big challenge in scRNA-seq analysis is that the gene expression is high dimensional and sparse, some of the data include more than 20,000 dimensions and 80% zero value. Embedding is one of the research problems to tackle this challenge. Projecting gene expression matrix from sparse space to low dimensional intense space helps to obtain meaningful insight from data. That is why several machine learning algorithms were proposed to reconstruct the gene expression matrix and then do some downstream analysis, including integration, imputation, prediction and cluster ([9],[8],[6],[10]). Because of the rapid progress in the single-cell research, there are many data sets generated from different experiments and groups. To improve the efficiency of the data and obtain more biological insight from different data set, data integration becomes a hot topic in scRNA-seq analysis ([11],[7],[12],[10]). These algorithms aim to project the points(cells) from different datasets to several clusters. In every cluster, they belong to the same cell type. However, this approach has drawbacks. Firstly, this approach can only integrate the datasets from the same species, because different species have different taxonomy. Secondly, when the dimensionality of the gene expression matrix is high, these algorithms would lose much information and the embedding from the data would lose much biological meaning.

Cross-species analysis, as a part of data integration, helps to reveal the evolutionary and developmental relationships between cell types across species. Many tools currently exist for analyzing single-cell data and identifying cell types. However, cross-species comparisons are complicated by many biological and technical factors [13]. There are several research try to align the cross-species data sets ([14],[10]). Nevertheless, they are either based on a biological process rather than data or project the data points(cells) to several clusters that contains the same cell type in each cluster.

The scRNA-seq expression data is sparse and has a large scale, which is similar to language data. And in natural language processing, Neural machine translation (NMT) [15] can be seem as embedding and transformation process, which is a kind of general data integration. NMT has witnessed rapid progress in recent years, from novel model structure developments [16] to achieving performance comparable to humans [17]. At the same time, unsupervised NMT, i.e., translate language without parallel data [18] has also witnessed maturation recently. To illustrate, they first train the language model for source and target language separately, then set the dictionary obtains from training Generative Adversarial Networks (GANs) [19] as initialization of the translator, finally train the translator via back-translation technique [20].

In summary, this paper makes the following main contributions:

1. We propose a pipeline that uses Wasserstein distance to compute the similarity of different cell types in different species.
2. The model we proposed can reveal biological insights better than any other embedding models used in scRNA-seq analysis, including autoencoder and variational autoencoder. Compare with the ground truth of taxonomy results from biologists [3], our model can obtain similar results in a data-driven approach.
3. We present a novel perspective to integrate scRNA-seq expression data.
4. We demonstrate that Transformers [21] can be used for general sparse and high-dimensionality data by visualizing the embedding results.

## 2 Background

### 2.1 Transformer

The Transformer architecture is usually developed by stacking Transformer layers [21]. A Transformer layer operates on a sequence of vectors and outputs a new sequence of the same shape. The computation inside a layer is decomposed into two steps: the vectors first pass through a (multi-head) self-attention sub-layer and the output will be further put into a position-wise feed-forward network sub-layer. Residual connection [22] and layer normalization [23] are employed for both sub-layers.

#### Self-attention sub-layer

The attention mechanism can be formulated as querying a dictionary with key-value pairs [21], e.g., 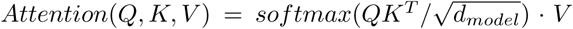, where *d*_*model*_ is the dimension of the hidden representation and *Q*(Query), *K*(Key), *V* (Value) are specified as the hidden representations of the previous layer in the so-called *self-attention* sub-layers in the Transformer architecture. The multi-head variant of attention allows the model to jointly attend to information from different representation subspace, and is define as

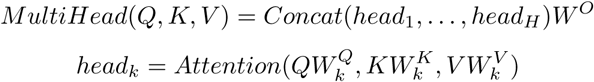

where 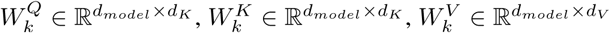, and 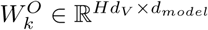 are project parameter matrices, *H* is the number of heads, and *d*_*K*_ and *d*_*V*_ are the dimension of Key and Value.

### 2.2 Generative Adversarial Networks

Generative Adversarial Networks(GANs) [19] usually contain generator network and discriminator network. In GANs, the generator network operates on a vector from source space and outputs a fake vector, and the discriminator network inputs the fake vector and a vector from the target space, then output the predicted label of the vector. In every training epoch, we first train the generator and then train the discriminator. In this process, we used adversarial loss that the generator tries to minimize the following function while the discriminator tries to maximize:

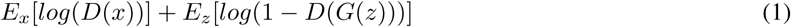

where *D*(*x*) is the discriminator’s estimate of the probability that real data instance *x* is from target space, *E*(*x*) is the expected value over all real data instances, *G*(*z*) is the generator’s output when given input *z*.

### 2.3 UMAP

UMAP (Uniform Manifold Approximation and Projection) [24] is a novel manifold learning technique for dimension reduction. UMAP is constructed from a theoretical framework based in Riemannian geometry and algebraic topology. The result is a practical scalable algorithm that applies to real-world data. The UMAP algorithm is competitive with t-SNE [25] for visualization quality, and arguably preserves more of the global structure with superior run time performance.

## 3 Model

In this paper, we propose two main models (i)An embedding model based on Transformers that can embed the high-dimensionality expression data. (ii)An algorithm that can measure the similarity of the cells in the embedding space we get from (i).

### 3.1 Our proposed architecture for embedding

We extend BERT(Bidirectional Encoder Representations from Transformers) [26] towards general sparse data sets and demonstrate the biological meaning of the embedding. Our model contains three main parts: linear transformation, transformer encoder and transformer decoder.

As shown in Figure 1, suppose we have dataset *X*_*A*_ ∈ ℝ^*a*×*n*^ and dataset *X*_*B*_ ∈ ℝ^*b*×*m*^, where *a* the amount of cells in the dataset *X*_*A*_, *n* is the amount of genes in the dataset *X*_*A*_, *b* the amount of cells in the dataset *X*_*B*_, *m* is the amount of genes in the dataset *X*_*B*_, the values of the matrix represent gene expression values for each cells. Notice that the expression matrix is sparse(typically over 80% of the values are zero).

**Figure 1:**
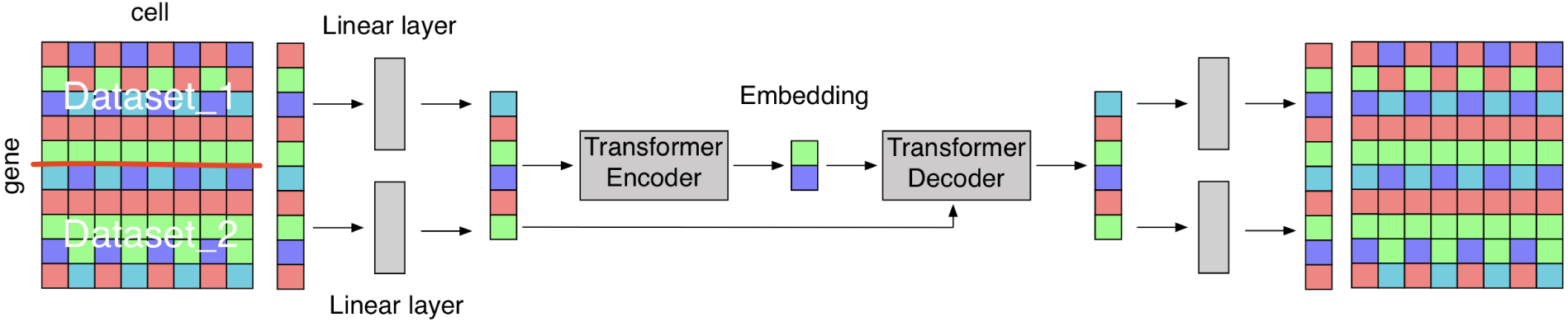
Overview of the model architecture

Through the linear layer *W*_*a*_ and *W*_*b*_, we can obtain the lossless intense representation of the expression matrix(since we can reconstruct the raw expression matrix by simple inverse transformation). e.g, the lossless intense representation of *X*_*A*_ and *X*_*B*_ are 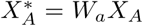 and 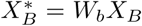. [26] shows that randomly mask some of the input and impute back is an efficient way to train a large model in an unsupervised approach. Then we randomly mask 10% of the data, which is a method to make the model more robust.

After linear layer, let *C*_*A*_ = *S*_*A*_*W*_*A*_ ∈ ℝ^1×*n*^ and *C*_*B*_ = *S*_*B*_*W*_*B*_ ∈ ℝ^1×*m*^ be the trainable source embedding that denote the source of the data, where *S*_*A*_ and *S*_*B*_ are one-hot vectors that denote the dataset. The input of the Transformer Encoder can be calculate as 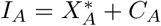 and 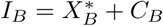

Then we can get the embeddings *E*_*A*_ and *E*_*B*_ from Transformer Encoder

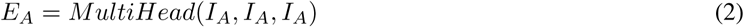

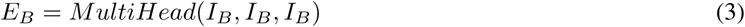

Inspired by the BERT [26], we utilize Attention mechanism to merge the information after Transformer Encoder

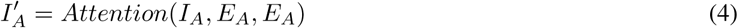

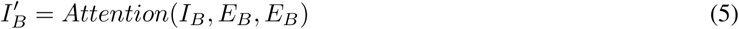

And the output of Transformer Decoder can be calculated as

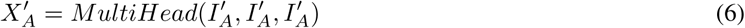

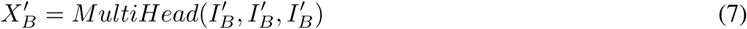

Finally, we use Mean Square Error 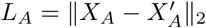 and 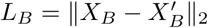 as loss function for reconstruction. Since the intense representation is lossless transformation of the raw data, it is equivalent to reconstruct data in the raw space.

### 3.2 scRNA-seq alignment

#### An Assumption

In this part, we assume that the cells in different experiments have similar functions and the functions can be determined by genes expression. To illustrate, in the language translation problem, we can assume that the embedding of English and German have a similar structure. Since we live in the same world, different languages can be seen as the different representations of the same things. The difference is that in scRNA-seq, the function of cells might be much different, and some datasets might contain the cells that are not included in another data set. Based on this assumption, although the manifold shape of the different datasets in the embedding space is similar, they can be aligned through some mathematical transformation.

Our model focuses on learning an alignment between two sets such that the manifold shape is similar in the shared embedding space. In this paper, we use the adversarial criterion to measure the similarity, which is denoted as ‖ · ‖_*D*_. We denote the source dataset as *𝒳* and the target matrix as *𝒴*, what we want to do is to find a transformation *W*^*^ such that

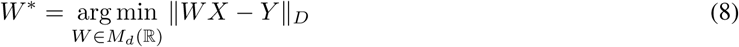

where *d* is the dimensionality of the embeddings, *M*_*d*_(ℝ) is the space of *d* × *d* matrices of real numbers, *X* is matrix of size *d* × *n* sampled from *𝒳, Y* is matrix of size *d* × *n* sampled from *𝒴*

In practice, [27] obtained better results on the word translation task using a simple linear mapping, and did not observe any improvement when using more advanced strategies like multilayer neural networks. And [18] construct linear language mapping through adversarial training, which demonstrates the effectiveness of learning linear mapping by GANs. So we use linear transformation to align the data and use GANs as the training framework.

Let *X* = *x*_1_, …, *x*_*n*_ and *Y* = *y*_1_, …, *y*_*n*_ be two sets sampled from *𝒳* and *𝒴*. A model is trained to discriminate between 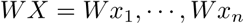 and *Y* = *y*_1_, …, *y*_*n*_. We call this model the discriminator. *W* is trained to prevent the discriminator from making accurate predictions. As a result, this is a two-player game, where the discriminator aims at maximizing its ability to identify the origin of an embedding, and *W* aims at preventing the discriminator from doing so by making *WX* and *Y* as similar as possible. The loss function fo discriminator is

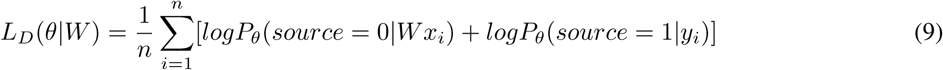

where *θ*_*D*_ is the parameter of discriminator, *source* = 0 means the data point is from source dataset and *source* = 1 means the data point is from target dataset

The loss function for alignment generator is

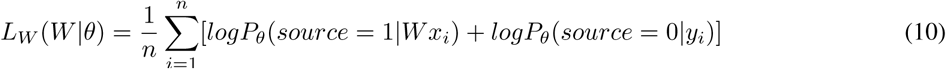

To train this model, we follow the standard training process of GANs [19], i.e for every training epoch, the discriminator and the mapping matrix *W* are trained successively with stochastic gradient updates to respectively minimize *L*_*D*_ and *L*_*W*_.

## 4 Related work

### Encoder-decoder architecture

As a basic algorithm in deep learning, encoder-decoder has been widely used in the analysis of scRNA-seq data. SAVER-X [6] integrate the scRNA-seq expression data via autoencoder and statistical learning method. SAUCIE (Sparse Autoencoder for Clustering, Imputing, and Embedding) [9] first explore the multitask learning based on the embeddings of autoencoder. scGen [8] has tried to represent the sparse scRNA-seq data by some low-dimension probability distribution via variational autoencoder.

### canonical correlation analysis (CCA)

The most famous computational tools in bioinformatics is Seurat [4]. The latest version [10] supports integrating diverse modalities associated with single-cell sequencing datasets, which can be used to better understand cellular identity and function. The key algorithm of the latest version of Seurat is canonical correlation analysis. [28] demonstrates that CCA can effectively capture correlated gene modules that are present in both datasets.

### Out of sample and cross-species experiment

[6] shows that transfer learning helps to improve the performance in several tasks, such as imputation and integration. In [8] and [6], they had experimented on the out-of-sample sets, i.e evaluate the algorithm in a dataset that contain some types of the sample did not appear on the training set. [14] focuses on cross-species alignment based on a biological process and it shows that cross-species analysis is meaningful for bioinformatics.

## 5 Experiment

We performed an extensive open set of computational experiments on several brain data sets, where we demonstrate the visualization of the result and compare the predictive performance of our proposed method against SAUCIE [9] and PCA [29]. The most popular model in data integration is Seurat [10] and it does not support to merge the data set with different genes. However, in our data, the genes of human and mouse are totally different.

### 5.1 Data Sets

We mainly use the brain atlas dataset from human and mouse. Because both of them are from the nervous system and there are many research projects ([3],[30],[31]) in this field, we can evaluate our findings by comparing with those results. In addition, large-scale and high-dimensionality brain atlas dataset is complex enough for data-driven algorithms.

#### 5.1.1 Human Brain Atlas (HBA)

Human Brain Atlas (HBA) (https://portal.brain-map.org/atlases-and-data/rnaseq#Human_Cortex) include 50,000 genes of 25,000 cells for more than 10 cell types. Individual layers of cortex were dissected from brain tissues including the middle temporal gyrus (MTG), anterior cingulate gyrus (CgGr), primary visual cortex (V1C), primary motor cortex (M1C), primary somatosensory cortex (S1C) and primary auditory cortex (A1C). Based on this dataset, [31] found conserved cell types with divergent features in the human versus mouse cortex.

#### 5.1.2 Mouse Brain Atlas (MBA)

Mouse Brain Atlas (MBA) (http://mousebrain.org/) includes 25,000 genes of 160,000 cells for more than 16 cell types, which provides a clearer picture of cell diversity by region and a reference atlas for studying the mammalian nervous system. It contains cells from more parts of bodies. The cells in Human Brain Atlas are only from Central Nervous System (CNS), while the Mouse Brain Atlas contains cells from Central Nervous System (CNS), Peripheral Nervous System (PNS) and Enteric Nervous System (ENS).

### 5.2 Experimental Settings

In data preprocessing, we first filter the cells and genes using the python package scprep (https://github.com/KrishnaswamyLab/scprep). Secondly, we log-transformed gene expression profiles. For each data set, we then scaled it to [−50, 50]. Finally, for each data set, we project it to 1200 dimensions as intense representation using linear Principle Component Analysis algorithm.

As for the hyperparameter, (i) in the embedding stage, the batch size is 1024, the learning rate is 0.0003, the embedding size for Transformers is 1024, the total amount of parameters in the embedding model is 300 million. (ii) In the alignment stage, we choose one layer linear networks without activation function as the generator and Multi-Layer Perceptron as the discriminator, the learning rate is 0.00003, the batch size is 1024.

In visualization, we first operate the 1024-dimensionality data through PCA and outputs 50-dimensionality data. Then input it to UMAP and get 10-dimensionality data. Finally, we choose the first two dimensions of data for visualization.

We implemented our models in PyTorch [32] using Google Cloud TPU-v3-8 with 128GB memory and it took four hours to get our results.

### 5.3 Evaluation Measure

We used Wasserstein distance to measure the similarity in geometry. Intuitively, if each distribution is viewed as a unit amount of “dirt” piled on *M*, the metric is the minimum “cost” of turning one pile into the other, which is assumed to be the amount of dirt that needs to be moved times the mean distance it has to be moved. The Wasserstein distance between distribution *u* and *v* is:

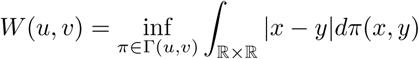

where Γ(*u, v*) is the set of distributions on ℝ × ℝ whose marginals are *u* and *v* on the first and second factors respectively.

To evaluate the biological significance of the embeddings, we first calculate the mean points of every cell type class and then used the Hierarchical Cluster [33] algorithm to reconstruct the taxonomy of the MBA data set. Then we calculate the accuracy level by level and compare our models with other methods.

### 5.4 Experimental Result

In the embedding task, we compared several machine learning algorithms, namely, multitask SAUCIE [9], PCA and our Transformers based model on two data sets. The visualization of embedding results is Figure 3.

**Figure 2:**
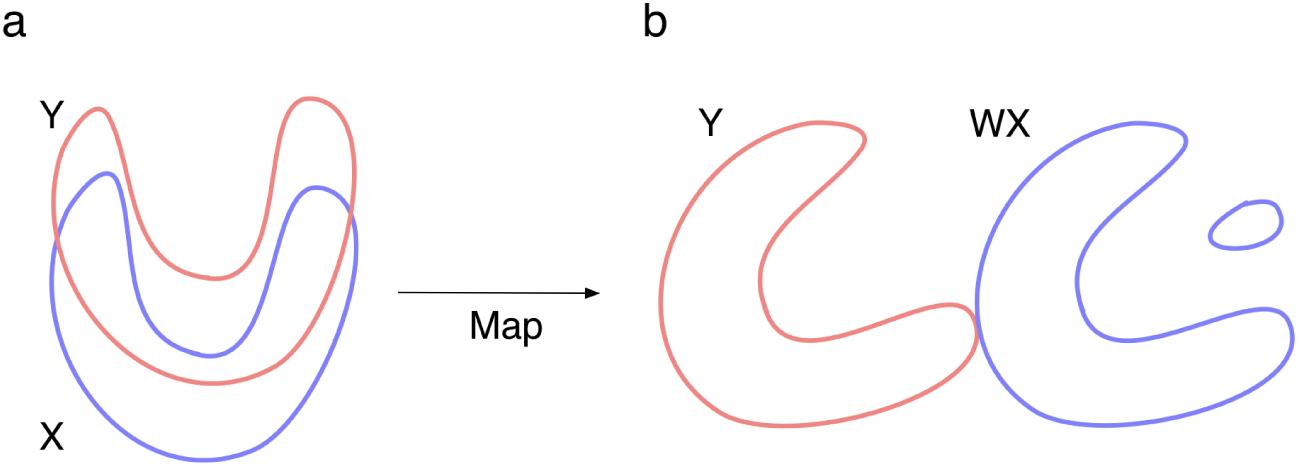
Overview of mapping. (**a**) Dataset X and dataset Y in the embedding space. (**b**) After mapping, assume that dataset X contains more cell types, dataset X would separate into two parts, one of them is similar to dataset Y. And the other part contains the cells that very different from all the cells in dataset Y.

**Figure 3:**
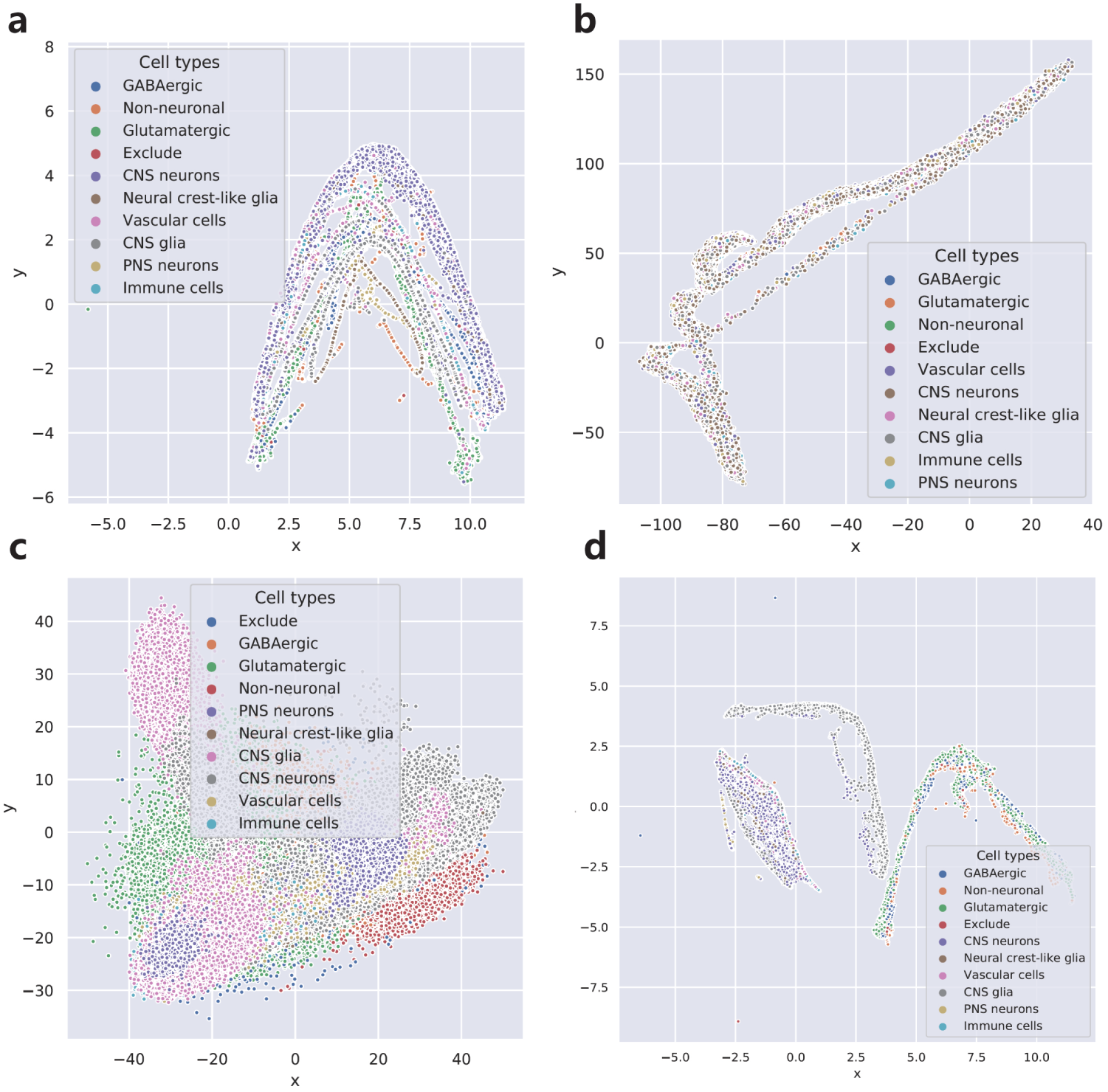
Main result of visualization. In the legend, the top 4 classes (*GABAergic, Non-neuronal, Glutamatergic, Exclude*) belong to HBA dataset, others are from MBA data set. (**a**)Visualization of embeddings from our model. (**b**) Visualization of embeddings from PCA. (**c**)Visualization of embeddings from SAUCIE[9]. (**d**)Visualization of aligned data from our model.The left two parts belongs to mouse and the right part belongs to human.

Figure 3 indicates that the embeddings of our models reserved biological knowledge better than the other two models. When we visualize the 1024-dimension embeddings in two-dimension space, our model (Figure 3 (**a**)) verified that since the two data sets (HBA and MBA) are operated through the same Transformer encoder and decoder, the embeddings of them are similar in the manifold shape. Besides, since we add source embedding to both data set before inputting into Transformer Encoder, the HBA data set and MBA data set can be separated. It also demonstrates that for each cell type, the distribution of cells in the embedding space is continuous and separated. As for the embedding of SAUCIE (Figure 3 (**b**)), the embedding space seems messy and highly compressed. The embedding of the PCA (Figure 3 (**c**)) is not as good as ours, it can neither discriminate different species nor cell types.

We observed that our alignment result (Figure 3 (**d**)) make biological sense. In the Method part of this paper, we expected that after our alignment, the different parts of the data would be separated from the main manifold. Interestingly, we found similar property in our result. The left part of data points is from MBA data set, which contains the cells from *CNS, PNS* and *ENS* while the right part of the data only from *CNS*. And the right part of the MBA data set, which is from *CNS*, is similar to the HBA data set.

We also observed that our result of hierarchical cluster (Figure 4 (**a**)) is very similar to the true taxonomy of MBA data set (Figure 4 (**b**)). In the top-level, our model separate into two parts(red and green branches), which correspond to Neurons and Non-neurons. We can consider such a problem: in the training stage, we do not know the labels of neurons/non-neurons, but we know the cell types for each sample, the task is to predict neurons/non-neurons for each sample.

**Figure 4:**
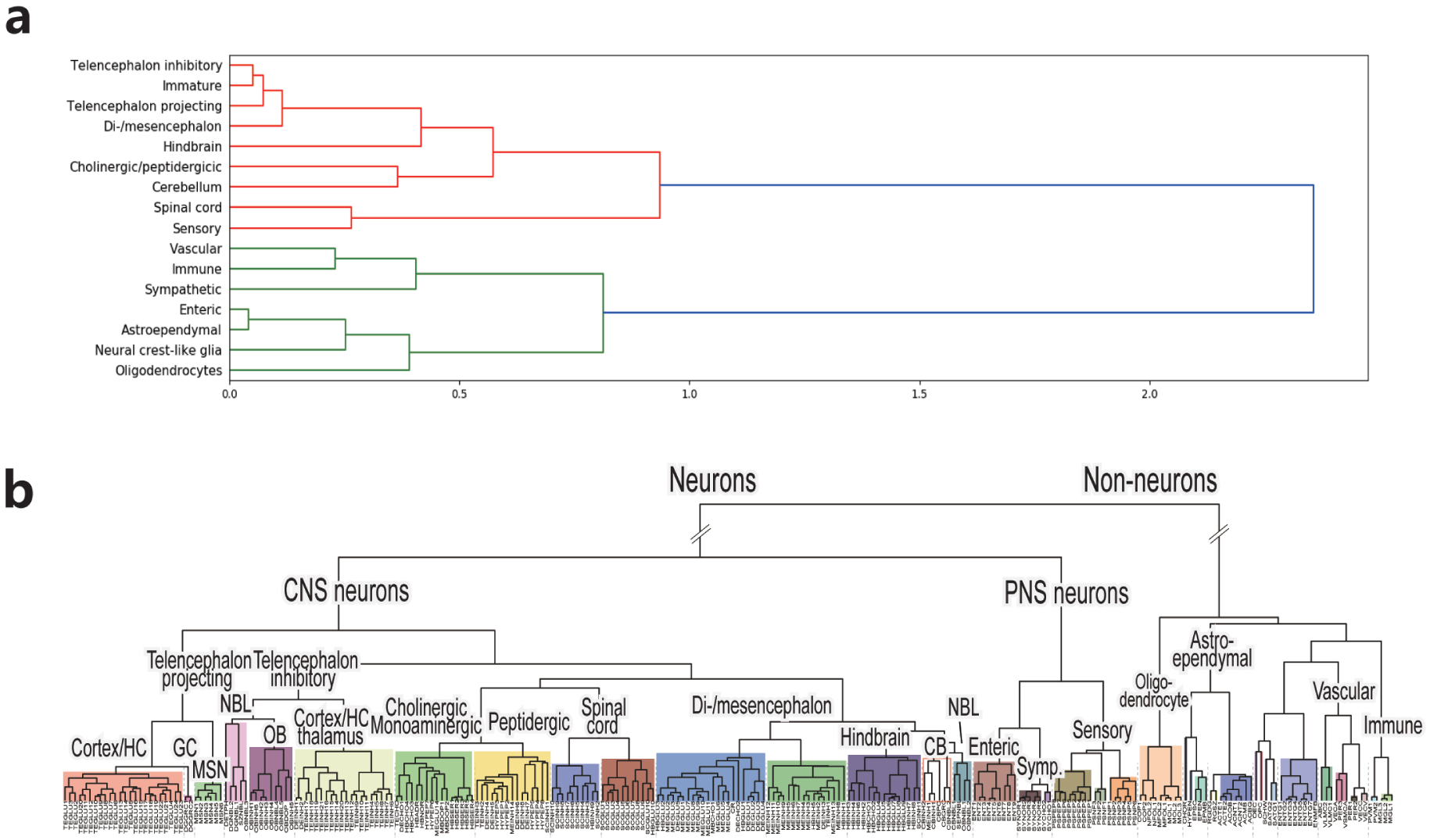
(**a**)Taxonomy generated from embedding space of our models. (**b**)Ground truth taxonomy from [3]

Our result of this unsupervised classification problem is good, 13 out of 15 cell types are right and the accuracy reaches 86.7% in the first level. The only two types get wrong classification results are *Sypathetic* and *Enteric*. But according to Figure 4 (**b**), the scale of these two classes is relatively small, so this error may be caused by the unbalanced data set. However, for the relatively large sets of cell types that are fully trained, such as *Oligodendrocytes, Di-/mesencephalon* and *Cholinergic/peptidergicic*, the position in taxonomy is very accurate.

We compare the result with SAUCIE and PCA in Table 1. We can not obtain the results of Seurat since it does not support to merge the data sets with different genes. In SAUCIE, they directly compress the data from high-dimension space to 2-dimension space. It is probable that the embedding is highly compressive, so its hierarchical information can not be discovered by hierarchical cluster directly. When clustered the data into neuron/non-neuron at the top level, the result is good. But when it goes the second level to discriminate CNS neurons from PNS neurons, the results got worse. In conclusion, our models show that Transformers architecture reserve the biological information well when it was used to embed the scRNA-seq expression matrix from high dimensional space to relatively low-dimension space.

**Table 1:** Cluster accuracy.

| MODELS | LEVEL-1 | LEVEL-2 |
| --- | --- | --- |
| SAUCIE[9] | 66.7% | 53.3% |
| PCA[29] | 80.0% | 60.0% |
| SEURAT[10] | - | - |
| **OURS** | **86.7%** | **66.7%** |

To quantify the property we found that our model reserved biological knowledge well, we use Wasserstein distance among each distribution of cell types to quantify the similarity in geometry. In practice, we normalize every cell-type cluster and then calculate the affinity matrix for the embeddings of MBA data set, i.e, compute per-pair of Wasserstein distance for each cell types[3] in MBA. Obviously, this affinity matrix (Figure 5) is symmetrical and the values in diagonal is zero. We observed that the values in the affinity matrix match the relationship shown in the Figure 4 (**b**), i.e, if the distance of two types in the Affinity matrix is large, the distance in the taxonomy would also be large. For example, the distance between neurons and non-neurons is relatively large in the affinity matrix. This findings have these significance: 1) It somehow illustrated why our models can capture the taxonomy information, i.e, manifold shape, distribution and variance. 2) Wasserstein distance, which is usually used in quantify the distance of two distribution, provides a good way to quantify the distance of different cell types in gene level. Because it is highly correlated with the distance in taxonomy.

**Figure 5:**
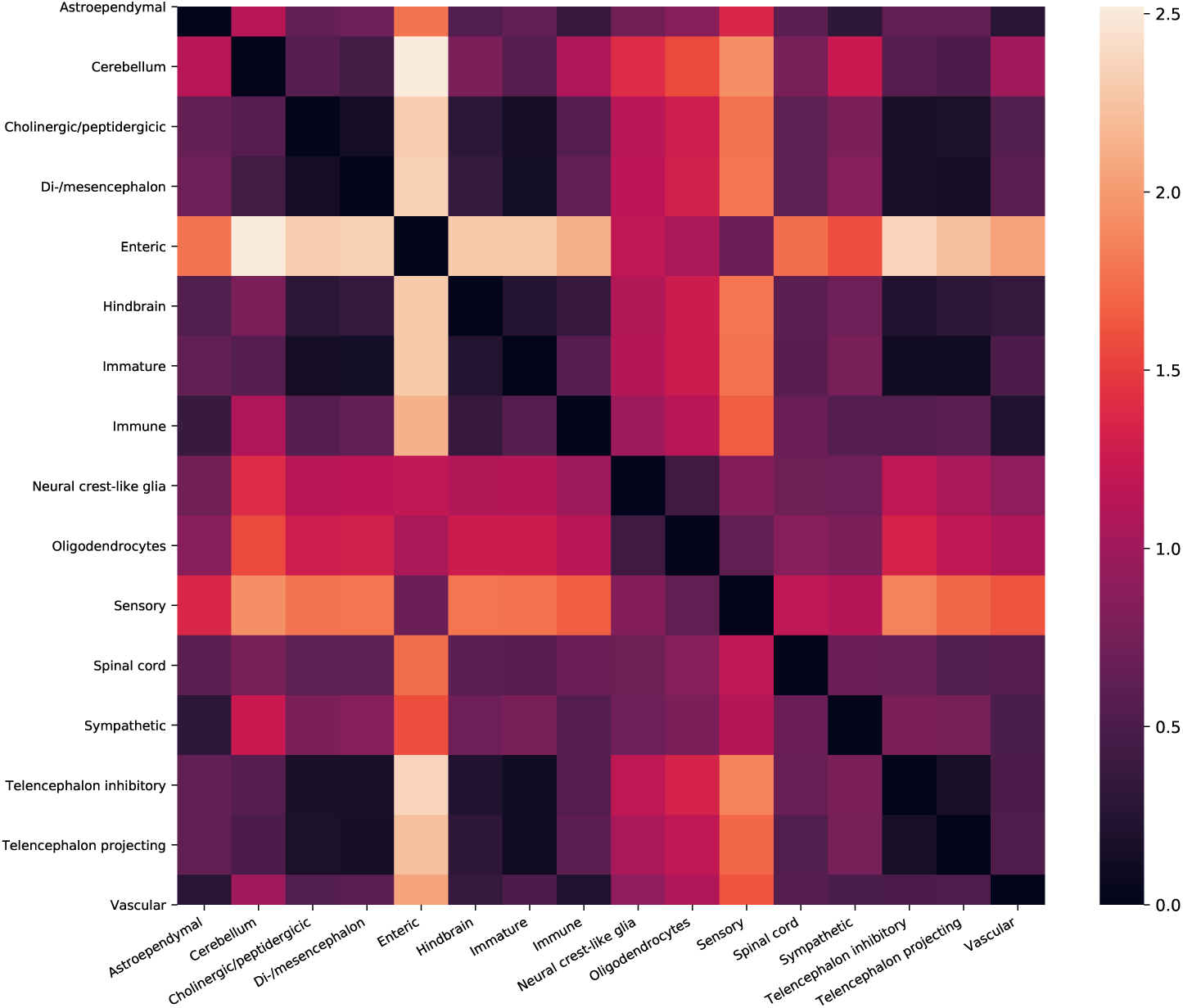
Affinity matrix.

In summery, Figure 3 shows our embeddings intuitively and aligned the data from different sets in a novel way. Figure 4 and Figure 5 demonstrates that our models can reserve the biological knowledge in both position and the manifold shape.

## 6 Conclusions

Embeddings and alignment for large-scale scRNA-seq data are quite important to better understand the relationship of cells across cell types and species. In this study, we extended BERT [26] towards general sparse data sets and utilized GANs [19] with linear generator to align the data in the manifold, which is know to obtain better mapping results [27].

To evaluate the models we proposed(Figure 1 and Figure 2), we used gene expression profiles from Human Brain Atlas and Mouse Brain Atlas [3].

We visualize the embedding and alignment results of our models and compared the results against PCA [29] and SAUCIE[9]. Our models obtain the best results in manifold analysis. Then we quantify and verify that the embeddings from our models can reserve the biological knowledge not only in the position of the points, but also in the manifold shape of the distribution.

For now, our models provides a pipeline for research on the similarity based on the Wasserstein distance in the embedding space. And our experiments demonstrate the power of large model(Transformers) and big data(Human and mouse brain atlas) for the analysis of scRNA-seq expression data.

We envision extending our models towards a new pipeline for preprocessing large-scale scRNA-seq data sets in the future. In this study, we trained shared Transformer Encoder and Transformer Decoder for two cross-species brain data sets with different gene lists. If we trained more data sets together for embedding, it is possible to obtain better results in many tasks in bioinformatics, just as what BERT has done in natural language processing.

